# Granulocyte-Colony Stimulating Factor reduces cocaine-seeking and downregulates glutamatergic synaptic proteins in medial prefrontal cortex

**DOI:** 10.1101/2020.02.17.949990

**Authors:** Rebecca S. Hofford, Tanner J. Euston, Rashaun S. Wilson, Katherine R. Meckel, Emily G. Peck, Arthur Godino, Joseph A. Landry, Erin S. Calipari, TuKiet T. Lam, Drew D. Kiraly

**Author notes:** Corresponding author: Drew D. Kiraly, 1 Gustave L Levy Pl – Box 1230, Icahn School of Medicine at Mount Sinai, New York, NY.

## Abstract

**Background:** Psychostimulant use disorder is a major public health issue, and despite the scope of the problem there are currently no FDA approved treatments. There would be tremendous utility in development of a treatment that could help patients both achieve and maintain abstinence. Previous work from our group has identified granulocyte-colony stimulating factor (G-CSF) as a neuroactive cytokine that alters behavioral response to cocaine, increases synaptic dopamine release, and enhances cognitive flexibility. Here, we investigate the role of G-CSF in affecting extinction and reinstatement of cocaine-seeking and perform detailed characterization of its proteomic effects in multiple limbic substructures.

**Methods:** Sprague-Dawley rats were injected with PBS or G-CSF during (1) extinction or (2) abstinence from cocaine self-administration, and drug seeking behavior was measured. Quantitative assessment of changes in the proteomic landscape in the nucleus accumbens (NAc) and medial prefrontal cortex (mPFC) were performed via data-independent acquisition (DIA) mass spectrometry analysis.

**Results:** Administration of G-CSF during extinction accelerated the rate of extinction, and administration during abstinence attenuated cue-induced cocaine-seeking. Analysis of global protein expression demonstrated that G-CSF regulated proteins primarily in mPFC that are critical to glutamate signaling and synapse maintenance.

**Conclusion:** Taken together, these findings support G-CSF as a viable translational research target with the potential to reduce drug craving or seeking behaviors. Importantly, recombinant G-CSF exists as an FDA-approved medication which may facilitate rapid clinical translation. Additionally, using cutting-edge multi-region discovery proteomics analyses, these studies identify a novel mechanism underlying G-CSF effects on behavioral plasticity.

## Introduction

Pathological substance use disorders (SUDs) are debilitating psychiatric conditions characterized by a pattern of escalating and dysregulated substance use followed by a chronic cycle of abstinence and relapse (American Psychiatric Association, 2013). In SUD patients, the propensity to relapse is a major impediment to successful treatment, and increases in cocaine craving following prolonged periods of abstinence have been seen in both human subjects and rodent models (Sinha, 2011; Parvaz et al., 2016). Despite the immense societal burden of illness, there are currently no FDA approved medications for the treatment of psychostimulant use disorder, and behavioral interventions alone are only marginally effective at ameliorating this disease (Penberthy et al., 2010). In recent years, there has been increased interest in non-traditional pharmacological approaches to treat cocaine addiction. There is growing evidence for neuroimmune interactions in multiple neuropsychiatric disorders including schizophrenia, depression, and addiction (Hodes et al., 2015; Lacagnina et al., 2017; Miller and Goldsmith, 2017; Hofford et al., 2019). This has led to an emerging body of research targeting neuroimmune interactions as possible translational research targets in addiction and other neuropsychiatric diseases.

Recently, we have identified granulocyte-colony stimulating factor (G-CSF) as a neuroactive cytokine with high relevance to SUD and motivation. While G-CSF was initially identified for its ability to mobilize stem cells from the bone marrow, numerous studies have found that centrally it functions as a trophic factor important for learning and memory (Diederich et al., 2009b), and it is protective in multiple models of neurological disease (Meuer et al., 2006; Ridwan et al., 2014). Recently, we found that G-CSF is increased in serum and brain following repeated cocaine, and levels of G-CSF positively correlated with cocaine intake (Calipari et al., 2018). Endogenous G-CSF signaling is necessary for the development of cocaine conditioned place preference (CPP) and enhancing levels of G-CSF through repeated systemic injections increases CPP and self-administration of low doses of cocaine (Calipari et al., 2018). In line with previous studies implicating G-CSF in learning and memory (Diederich et al., 2009a, 2009b), we also demonstrated that G-CSF enhanced behavioral flexibility in an operant reversal task (Kutlu et al., 2018). Together, this suggests that G-CSF might be influencing cocaine reward and reinforcement by affecting motivation for cocaine or learning processes associated with operant self-administration.

Two of the most common ways to assess relapse in preclinical models are reinstatement after extinction and re-exposure to drug cues after a period of abstinence (Grimm et al., 2001; Dong et al., 2017; Farrell et al., 2018). All models of relapse involve recall of previously learned response-outcome contingencies and require intact functioning of the mesolimbic dopaminergic system (Kalivas et al., 2005; Hyman et al., 2006; Farrell et al., 2018), parts of which are known to be affected by G-CSF (Mervosh et al., 2018; Brady et al., 2019). Active self-administration, extinction, and reinstatement all recruit NAc and medial prefrontal cortex (mPFC) but extinction and reinstatement more heavily rely upon mPFC (Kalivas and McFarland, 2003; Hyman et al., 2006; Peters et al., 2009), making these areas attractive targets of G-CSF action in cocaine reinforcement, extinction, and reinstatement.

Given the potential for G-CSF to alter the behavioral plasticity associated with drug-seeking, the current study measured the effect of G-CSF on both cocaine extinction and reinstatement. Since G-CSF potentiated cocaine reward in CPP, increased cocaine intake during self-administration (Calipari et al., 2018), and enhanced reversal learning (Kutlu et al., 2018), these experiments helped determine if G-CSF exerts its effects on motivation or learning processes. Finally, to explore the underlying molecular mechanisms of G-CSF, we performed comprehensive discovery proteomics analysis on the NAc and dorsal mPFC following cocaine reinstatement. The insights gained into the mechanism of action of G-CSF could support the repurposing of G-CSF or the design of more targeted therapeutics for psychostimulant use disorder.

## Methods and Materials

### Animals and Housing

Adult male Sprague-Dawley rats (ENVIGO-Harlan), weighing between 280 – 300 g were pair-housed in a room with controlled temperature and humidity on a 24 hr reverse light-dark cycle (lights on at 19:00). All rats had *ad libitum* access to food and water before the start of any experiments. All animal procedures were approved by the IACUC at the Icahn School of Medicine at Mount Sinai and all practices conformed to the “Guide for the Care and Use of Laboratory Animals” (National Research Council 2010). Separate rats were used in Experiment 1 and Experiment 2.

### Cocaine Self-Administration

Self-administration, extinction, and reinstatement occurred in standard two-lever operant boxes equipped with two retractable levers, a syringe pump, and two jewel lights placed above each lever (Med Associates Inc, St. Albans VT). Before surgery, rats were anesthetized with a ketamine/xylazine cocktail (100mg/kg / 10mg/kg) before placement of an indwelling catheter (Plastics One, Torrington, CT) into the right jugular vein. Rats from Experiments 1 and 2 were allowed to recover for 3 to 7 d before the start of self-administration. Sessions were initiated by the insertion of an active and inactive lever. Lever presses on the active lever were reinforced on a fixed ratio 1 schedule and resulted in a 5.9 s infusion of 0.8 mg/kg/infusion (0.1 ml) cocaine and concurrent illumination of both cue lights (Experiment 1) or the cue light above the active lever (Experiment 2). There was a signaled time-out of 20 sec; active lever presses during the time-out were recorded but did not result in cocaine infusion. Active self-administration sessions lasted 3 hr once daily for 10 d. Presses on the inactive lever had no programmed consequence. Position of the active lever was counterbalanced across rats. All rats were placed on food restriction starting the day before self-administration start and continuing throughout the entire study by administration of 18 g food / rat delivered after the end of the session.

### Extinction

Rats in Experiment 1 and Experiment 2 underwent extinction. Extinction sessions ran daily for 5 d starting the day after rats’ last self-administration session. Sessions were 3 hr and were initiated as described above. During this phase, active lever presses no longer had any programmed consequence.

### Abstinence and Reinstatement

Rats in Experiment 2 were left in their home cages for 7 d following extinction training. For cue-induced reinstatement, rats were returned to the operant boxes for 30 min where responding on the previously active lever caused illumination of the light cue but did not result in cocaine infusion. Rats were returned to the colony and remained in their home cage for another 4 d. For cocaine-primed reinstatement, rats were injected with 10 mg/kg cocaine (i.p.) immediately before placement into the operant boxes for 30 min where lever presses on the previously active lever had no programmed consequence.

### Drugs

Cocaine hydrochloride was provided by the NIDA drug supply program from National Institute on Drug Abuse and was diluted in saline. Rat granulocyte colony stimulating factor (G-CSF) was purchased from GenScript Corp (Piscataway NJ) and was diluted with sterile phosphate buffered saline (PBS) to a concentration of 50 µg/ml.

### Experimental Timeline

#### Experiment 1

After self-administration training, rats were divided into PBS and G-CSF groups, such that levels of responding during acquisition did not differ. Rats received either PBS or 50 µg/kg G-CSF (i.p.) once daily 30 mins prior to the start of each extinction session. Five rats failed to acquire cocaine self-administration and were dropped from the study before extinction due to patency loss for final subject sizes of PBS *n* = 6 and G-CSF *n* = 7.

#### Experiment 2

Rats were divided into PBS and G-CSF treatment groups at the end of extinction, such that levels of responding during acquisition and extinction did not differ between groups. Rats were injected once daily with PBS or 50 µg/kg G-CSF (i.p.) during abstinence and 30 mins prior to each reinstatement session. Seven rats were dropped from the study before extinction due to loss of patency or, in one instance, inconsistent cocaine intake over days for final subject sizes of PBS *n* = 8 and G-CSF *n* = 9.

### Tissue Collection

Rats from Experiment 2 were euthanized by rapid decapitation; NAc core and dorsal mPFC (consisting of anterior cingulate and prelimbic cortices) were extracted and flash frozen 30 min after the conclusion of the cocaine-primed reinstatement test.

### Sample Preparation for LC–MS/MS

NAc and mPFC tissues were lysed with a probe sonicator in solubilization buffer (8 M urea 0.4mM, ammonium bicarbonate pH 8). Lysate was centrifuged at max speed for 10 minutes at 10 °C in a tabletop centrifuge to pellet cellular debris. Supernatant containing proteins (50 µg) was placed into an Eppendorf tube, and the volume was adjusted to 100 µL with solubilization buffer. Proteins were reduced with 5 µL of 200 mM dithiothreitol (DTT) and incubated at 37 °C for 30 min. They were then alkylated with 5 µL of 500 mM iodoacetamide (IAM) and incubated in the dark at room temperature for 30 min. After diluting with water to bring urea concentration to 2 M, sequencing-grade trypsin (Promega, Madison, WI, USA) was added at a weight ratio of 1:20 (trypsin/protein) and incubated at 37 °C for 16 h. The digested samples were then acidified with 0.1% formic acid, desalted using C18 spin columns (The Nest Group, Inc., Southborough, MA, USA), and dried in a rotary evaporator. The samples were resuspended in 0.2% trifluoroacetic acid (TFA) and 2% acetonitrile (ACN) in water prior to LC–MS/MS analysis.

### Data-Independent Acquisition (DIA)

DIA LC–MS/MS was performed using a nanoACQUITY UPLC system (Waters Corporation, Milford, MA, USA) connected to an Orbitrap Fusion Tribrid (ThermoFisher Scientific, San Jose, CA, USA) mass spectrometer. After injection, the samples were loaded into a trapping column (nanoACQUITY UPLC Symmetry C18 Trap column, 180 µm × 20 mm) at a flow rate of 5 µL/min and separated with a C18 column (nanoACQUITY column Peptide BEH C18, 75 µm × 250 mm). The compositions of mobile phases A and B were 0.1% formic acid in water and 0.1% formic acid in ACN, respectively. The peptides were separated and eluted with a gradient extending from 6% to 35% mobile phase B in 90 min and then to 85% mobile phase B in additional 15 min at a flow rate of 300 nL/min and a column temperature of 37 °C. Column regeneration and up to three blank injections were carried out in between all sample injections. The data were acquired with the mass spectrometer operating in a data-independent mode with an isolation window width of 25 m/z. The full scan was performed in the range of 400–1,000 m/z with “Use Quadrupole Isolation” enabled at an Orbitrap resolution of 120,000 at 200 m/z and automatic gain control (AGC) target value of 4 × 105. Fragment ions from each peptide MS2 were generated in the C-trap with higher-energy collision dissociation (HCD) at a collision energy of 28% and detected in the Orbitrap at a resolution of 60,000.

DIA spectra were searched against a Rattus norvegicus brain proteome fractionated spectral library generated from DDA LC MS/MS spectra (collected from the same Orbitrap Fusion mass spectrometer) using Scaffold DIA software v. 1.1.1 (Proteome Software, Portland, OR, USA). Within Scaffold DIA, raw files were first converted to the mzML format using ProteoWizard v. 3.0.11748. The samples were then aligned by retention time and individually searched with a mass tolerance of 10 ppm and a fragment mass tolerance of 10 ppm. The data acquisition type was set to “Non-Overlapping DIA”, and the maximum missed cleavages was set to 2. Fixed modifications included carbamidomethylation of cysteine residues (+57.02). Dynamic modifications included phosphorylation of serine, threonine, and tyrosine (+79.96), deamination of asparagine and glutamine (+0.98), oxidation of methionine and proline (+15.99), and acetylation of lysine (+42.01). Peptides with charge states between 2 and 4 and 6–30 amino acids in length were considered for quantitation, and the resulting peptides were filtered by Percolator v. 3.01 at a threshold FDR of 0.01. Peptide quantification was performed by EncyclopeDIA v. 0.6.12 (Searle et al., 2018), and six of the highest quality fragment ions were selected for quantitation. Proteins containing redundant peptides were grouped to satisfy the principles of parsimony, and proteins were filtered at a threshold of two peptides per protein and an FDR of 1%.

### Pathway analysis

Proteins were excluded from analysis if they were not detected in > 50% of all samples irrespective of treatment. Pairwise comparisons of the Log_10_ median intensity of every remaining protein and protein group were made using Scaffold DIA proteomics analysis software (http://www.proteomesoftware.com/products/dia/). Significantly upregulated downregulated proteins from each brain region were separately uploaded into the open source pathway analysis software package G:Profiler (Raudvere et al., 2019) (https://biit.cs.ut.ee/gprofiler/gost) to identify significantly enriched gene ontologies and KEGG pathways using an FDR corrected *p* < 0.05. Given that this software utilizes gene names for identifying pathways, all protein names were converted to gene names prior to pathway analysis using the Uniprot database (https://www.uniprot.org/uploadlists/). Tables listing all protein to gene name conversions in each region are available as **Tables S1-S2.** To generate **Fig. 5**, related pathway terms were reduced using Revigo (Supek et al., 2011) (http://revigo.irb.hr/) and portions of figure were created with BioRender.com. Upregulated and downregulated proteins from NAc and mPFC were separately uploaded into the STRING database (Szklarczyk et al., 2019) (https://string-db.org/) and enrichment of protein-protein interactions was assessed with the “multiple proteins” query using default settings. Statistical analyses for protein-protein interaction (PPI) enrichment, and number of predicted interactions per protein (average node degree), were performed utilizing the STRING analysis software. For visualization of protein-protein interaction networks disconnected nodes were removed from the image, and node size was adjusted to correspond to the −log(*p*) relative to PBS control.

**Figure 1.**
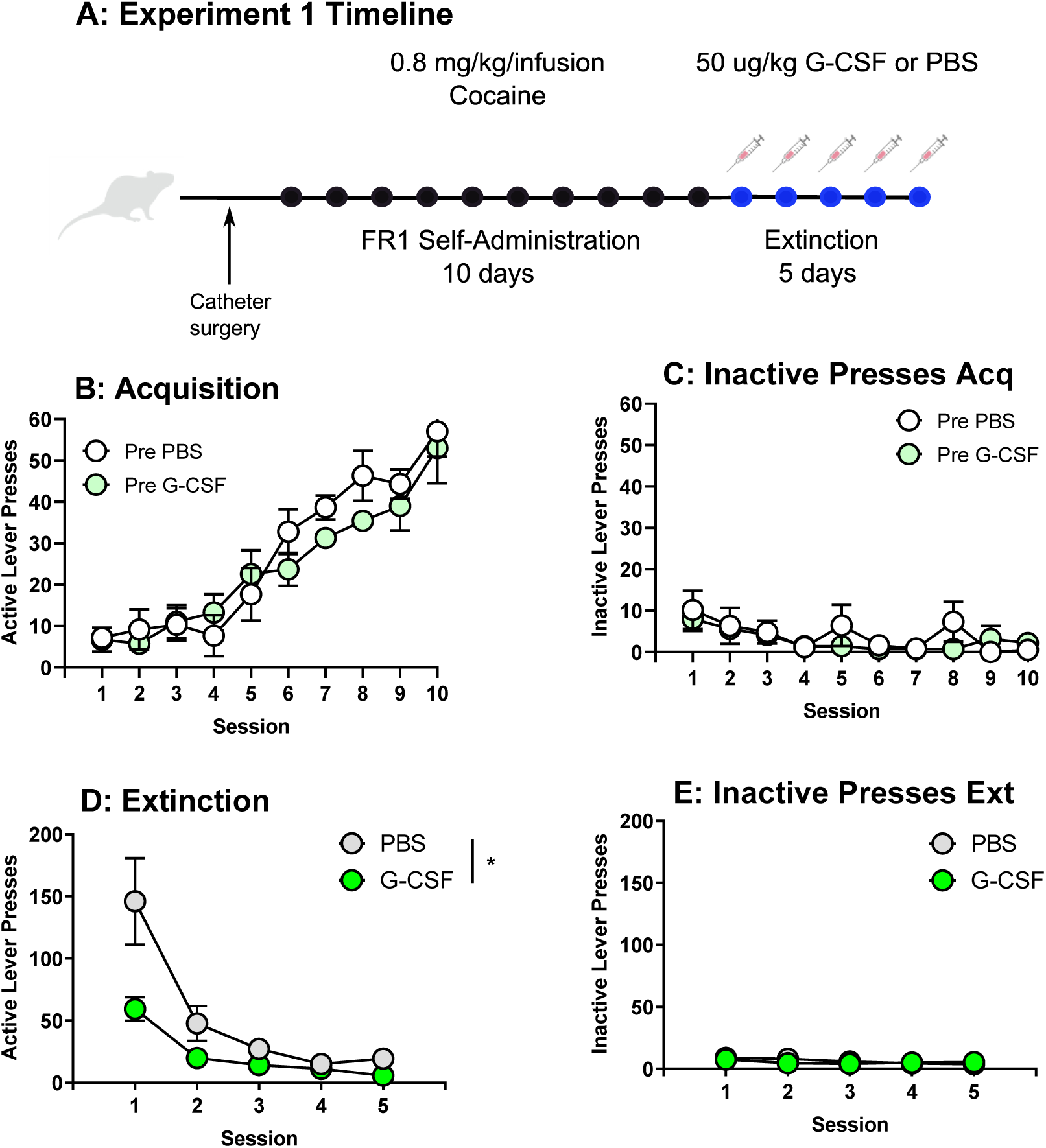
G-CSF accelerates extinction. (**A**) Schematic for Experiment 1. (**B**) Acquisition of active responding for cocaine (0.8 mg/kg/infusion, FR1) for rats later assigned to PBS or G-CSF groups showed no group differences (*F*_(1, 11)_ = 0.79, *p* > 0.05). (**C**) Pre-treatment groups also did not differ on inactive lever presses (*F*_(1,11)_ = 0.48, *p* > 0.05). (**D**) During extinction injections of G-CSF reduced lever pressing on the previously active lever (*F*_(1,11)_ = 5.37, *p* < 0.05) but did not affect rates of inactive lever pressing (**E** – *F*_(1, 11)_ = 0.10, *p* > 0.05). All data presented as means ± SEM, * *p* < 0.05, main treatment effect. Created with BioRender.com.

**Figure 2.**
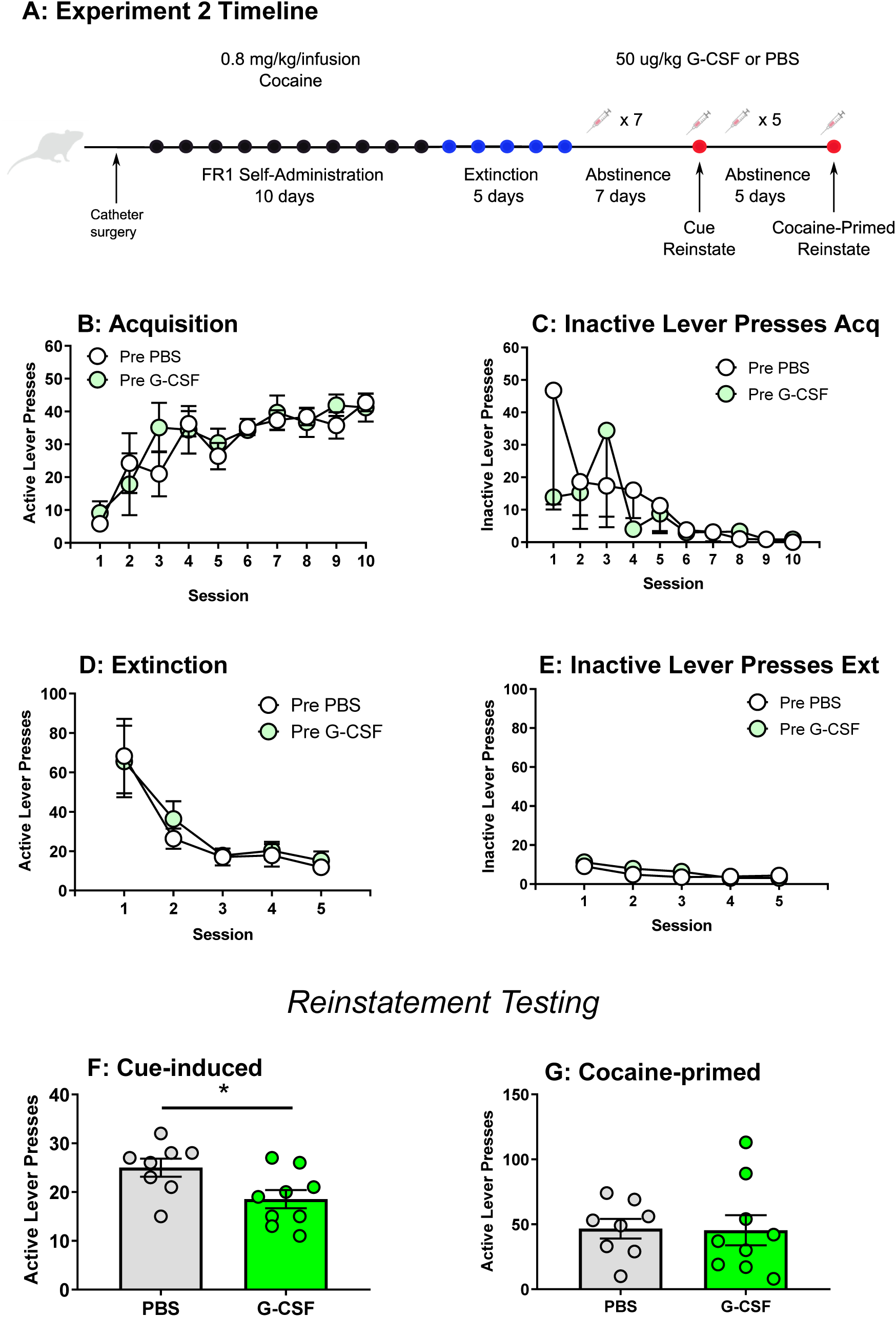
G-CSF reduces cue- but not cocaine-primed reinstatement when administered during abstinence. (**A**) Schematic for Experiment 2. (**B**) Rats later assigned to PBS or G-CSF groups did not differ in their active (*F*_(1, 15)_ = 0.21, *p* > 0.05) or (**C**) inactive pressing during acquisition of cocaine self-administration (0.8 mg/kg/infusion, FR1), but there was a main effect of session on active lever pressing (*F*_(3.84, 52.86)_ = 8.77, *p* < 0.0001). Additionally, pre-treatment groups did not differ in active (**D** – *F*_(1, 15)_ = 0.15, *p* > 0.05) lever pressing or (**E**) inactive lever presses during extinction, but there was a main effect of session on previously active lever pressing (*F*_(1.13, 13,53)_ = 15.77, *p* < 0.01). (**F**) G-CSF treatment during abstinence reduced responding for cue-induced reinstatement (*t*_(15)_ = 2.45, *p* < 0.05). However, there was no effect during cocaine-primed reinstatement (**G** – *t*_(15)_ = 0.08, *p* > 0.05) cocaine-primed reinstatement. All data presented as means ± SEM, * *p* < 0.05, main treatment effect. Created with BioRender.com.

**Figure 3.**
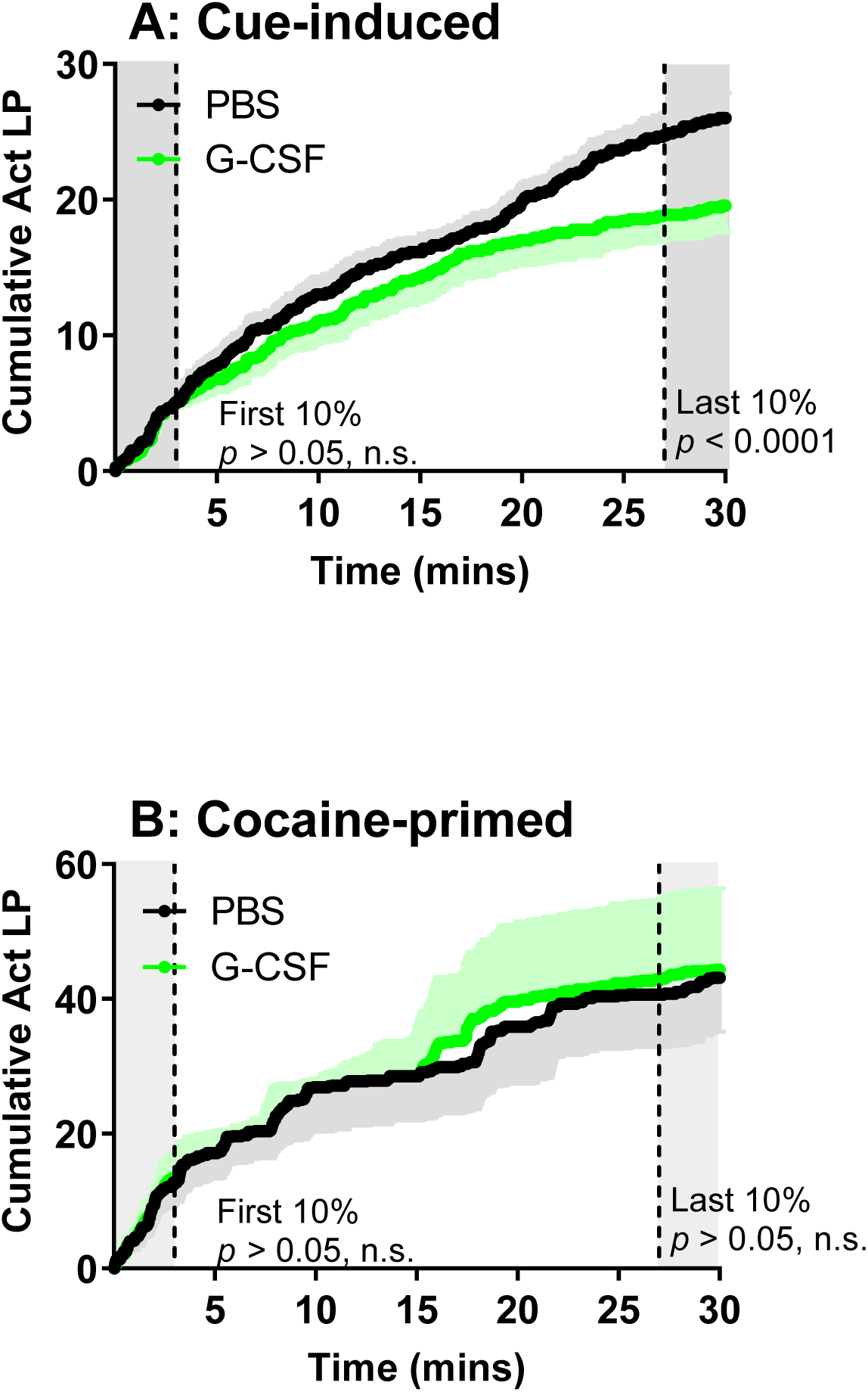
G-CSF treatment leads to updated response output over the course of cue-induced reinstatement session. (**A**) G-CSF and PBS treated rats did not differ in lever pressing within the first ten percent of the cue-induced reinstatement session (*t*_(15)_ = 1.79, *p* > 0.05), but significantly differed during the final 10% of the session (*t*_(15)_ = 16.25, *p* < 0.0001). (**B**) G-CSF and PBS treated rats did not differ in the first 10% of their cocaine-primed reinstatement session nor the final 10% (first 10%: *t*_(15)_ = 0.01, *p* > 0.05; final 10%: *t*_(15)_ = 0.92, *p* > 0.05). Black lines = PBS, green lines = G-CSF, black and green shading colors above and below indicate SEM.

**Figure 4.**
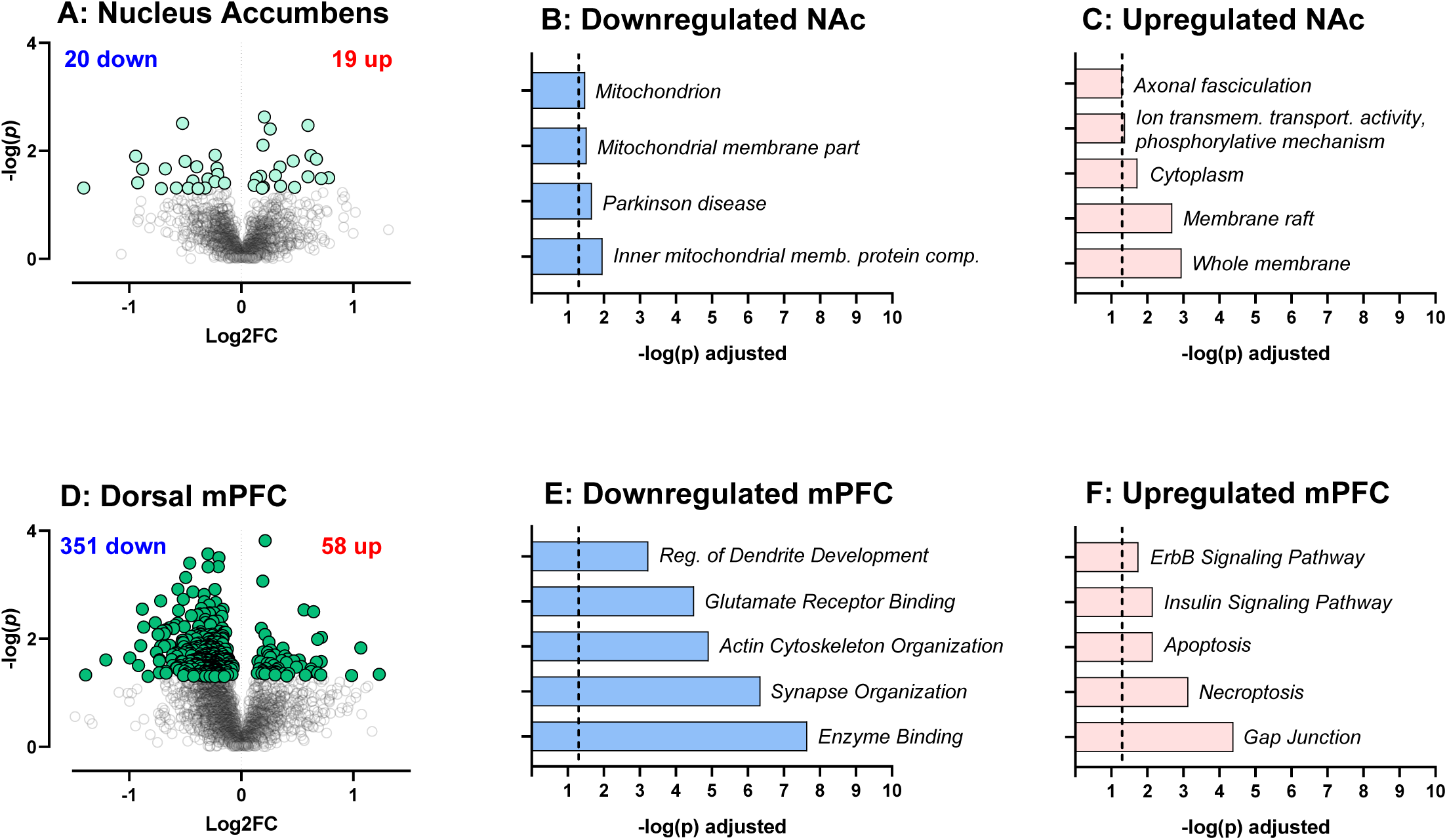
Proteomic effects of G-CSF on nucleus accumbens and dorsal medial prefrontal cortex. (**A**) Volcano plot depicting differential protein expression in the NAc between rats treated with G-CSF compared to PBS controls. (**B-C**) Significantly altered pathways in NAc identified from G:Profiler using significantly downregulated & upregulated inputs. (**D**) Volcano plot depicting differential protein expression between rats treated with G-CSF compared to PBS in mPFC. (**E-F**) Pathways identified from G:Profiler using significantly downregulated proteins (**E**) and significantly upregulated proteins (**F**) in mPFC. For **B**,**D**,**E**,**F** X-axis is adjusted −log p-value of the pathway; dashed line represents the level of significance (FDR-corrected *p* < 0.05).

**Figure 5.**
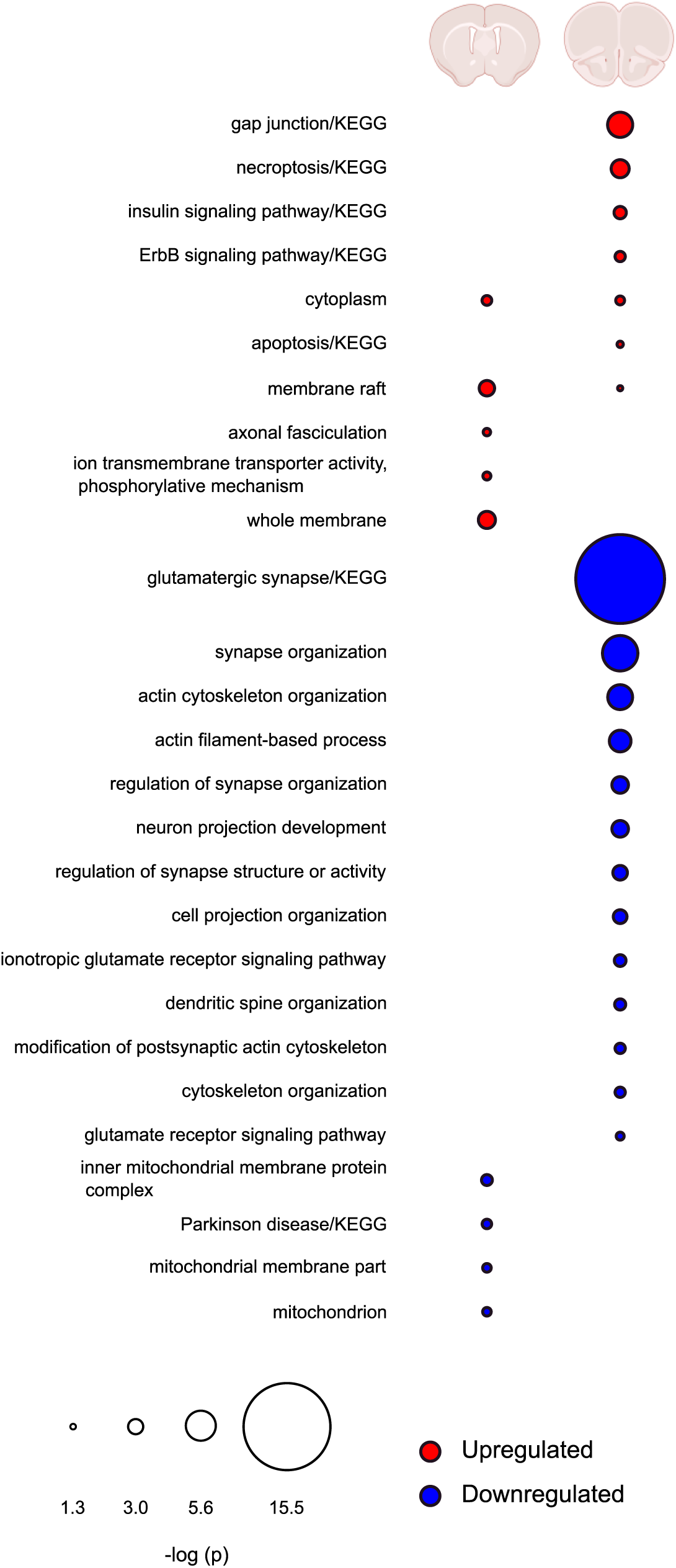
Comparison of regulated pathways in mPFC and NAc after G-CSF treatment. Comparison chart showing overlap of pathways found to be significantly enriched by G-CSF treatment in either NAc (**left**) or mPFC (**right**). Upregulated pathways are indicated in red, downregulated in blue. Radius of circles is the FDR-corrected −log *p* value of that pathway. Blank spaces for either brain region indicate no proteins from that pathway were significantly regulated by G-CSF in that region. Created with BioRender.com.

### Statistical analysis

For self-administration and extinction, data were analyzed separately using linear mixed effects with group (PBS or G-CSF) as a between-subjects fixed factor and session as a within-subject fixed factor. Significant interactions were probed with post-hoc analyses using Holm-Sidak. Lever pressing during cue- and cocaine-primed reinstatement were analyzed using independent samples t-tests. Greenhouse-Geisser correction was applied when appropriate. Time course analysis over the course of reinstatement were performed by independent samples t-tests of area under the curve between PBS treated and G-CSF treated rats during the first 3 min and the last 3 min. For proteomics, t-tests between PBS and G-CSF treated raw intensity values were conducted using Scaffold DIA software using an uncorrected *p* < 0.05.

## Results

### Experiment 1: G-CSF facilitates extinction of cocaine self-administration

Previous studies implicate G-CSF in enhancing cognitive flexibility on a reversal learning task (Kutlu et al., 2018). Given its ability to enhance motivated learning, we postulated that administration of G-CSF during extinction might hasten extinction learning. Rats were trained to self-administer cocaine and were split into two equally administering groups that would then be treated with G-CSF or PBS (**Fig. 1A**). No group differences in active (**Fig. 1B** – *F*_(1, 11)_ = 0.79, *p* > 0.05) or inactive (**Fig. 1C** – *F*_(1, 11)_ = 0.48, *p* > 0.05) lever pressing was found between rats that were later assigned into the PBS or G-CSF groups. As expected, only a main effect of session was found for active lever pressing during acquisition (**Fig. 1B** – *F*_(9, 97)_ = 33.52, *p* < 0.0001). However, during extinction, rats treated with G-CSF showed decreased responding on the previously active lever beginning on the first day of extinction (**Fig. 1D**). Throughout the sessions there was a main effect of treatment *F*_(1, 11)_ = 5.37, *p* < 0.05, as well as session *F*_(1.2, 11.13)_ = 23.4, *p* < 0.001, and a significant treatment x session interaction *F*_(4, 37)_ = 4.63, *p* < 0.01. However, post-hoc analysis comparing G-CSF and PBS treated active lever presses on each day did not yield any significant individual day effects (**Fig. 1D**). Importantly, treatment with G-CSF did not lead to any changes in responding on the inactive lever (**Fig. 1E** – *F*_(1, 11)_ = 0.10, *p* > 0.05). Additionally, to test if reduced extinction responding was due to a decrease in activity, we monitored locomotor activity within the operant chamber across days, and this was also not affected by treatment with G-CSF (**Fig. S1**).

### Experiment 2: G-CSF attenuates cue-but not cocaine-primed reinstatement

Given that G-CSF attenuated extinction responding beginning on the first day of training, we next investigated if G-CSF treatment during abstinence could reduce drug seeking. For experiment 2, a different set of animals were trained to self-administer cocaine on an FR1 schedule and then underwent five days of extinction prior to any treatments (**Fig. 2A**). Animals were divided into groups with equal levels of self-administration and inactive lever presses during the acquisition phase (**Fig. 2B,C** – active lever presses treatment effect: *F*_(1, 15)_ = 0.21, *p* > 0.05, active lever press session effect: *F*_(3.84, 52.86)_ = 8.77, *p* < 0.0001), as well as with equal responding on previously active and inactive levers during the extinction phase of training (**Fig. 2D,E**-formerly active lever presses treatment effect: *F*_(1, 15)_ = 0.15, *p* > 0.05; formerly active lever presses session effect: *F*_(1.13, 13,53)_ = 15.77, *p* < 0.01). When drug seeking was measured, daily injections of G-CSF during abstinence significantly attenuated cue-induced reinstatement (**Fig. 2F** -*t*_(15)_ = 2.45, *p* < 0.05). When cocaine-primed reinstatement was assessed after five more days of abstinence and G-CSF injections, there was no effect of G-CSF (**Fig. 2G** -*t*_(15)_ = 0.08, *p* > 0.05). G-CSF did not affect inactive lever presses during either cue or cocaine-primed reinstatement (data not shown).

### G-CSF reduces lever pressing late vs early within the cue-induced reinstatement session

Our previously published data on reversal learning (Kutlu et al., 2018) and the current data from the extinction and reinstatement experiments all suggested that G-CSF might be enhancing the ability of animals to rapidly update their response contingencies. To examine this in more detail, we analyzed cumulative responding on the previously active lever within each reinstatement session. As shown in **Fig. 2F**, G-CSF treated rats pressed the previously active lever less than control rats. However, within the first 10% (3 minutes) of the session, there was no difference in cumulative active lever presses between groups as assessed by comparison of area under the curve (**Fig. 3A** -*t*_(15)_ = 1.79, *p* > 0.05) indicating that G-CSF rats are sampling the active lever at the same rate as controls early in the session. However, by the last 10% of the session the G-CSF treated animals have decreased pressing on the previously active lever to levels below those of the control animals (*t*_(15)_ = 16.25, *p* < 0.0001). This suggests that increased levels of G-CSF might lead to rapid adjustment of behavior in response to altered reward availability over the course of a single session. In contrast, G-CSF did not significantly affect the rate of responding across a cocaine-primed reinstatement session (**Fig. 3B** -first 10%: *t*_(15)_ = 0.01, *p* > 0.05; final 10%: *t*_(15)_ = 0.92, *p* > 0.05)

### G-CSF given during abstinence alters the proteomic landscape of the mPFC but only minimally affects protein expression in the NAc

While evidence suggests that G-CSF alters dopamine dynamics in the NAc (Kutlu et al., 2018; Brady et al., 2019), the full mechanisms by which it alters behavioral plasticity are currently unknown. To gain insight into the neural processes underlying the effect of G-CSF on attenuated cocaine-seeking, a data-independent acquisition (DIA) discovery proteomic analysis was performed on two areas of the corticolimbic reward circuit in G-CSF and PBS treated rats from experiment 2. This method allows for unbiased detection of quantitative differences between treatment groups with improved coverage of the proteome compared to more conventional data dependent acquisition methods (Wilson et al., 2019). Both NAc and dorsal mPFC from rats administered G-CSF or PBS during abstinence and reinstatement (Experiment 2) underwent proteomic analysis. G-CSF treatment resulted in differential expression of 39 proteins in NAc (20 upregulated and 19 downregulated, **Fig. 4A, Table S3**). Pathway analysis of significantly up and down regulated proteins showed very few significant ontologies, with membrane rafts and membrane proteins being the most significant pathways (**Fig. 4B,C**). In contrast, analysis of G-CSF effects in the mPFC led to robust regulation of many proteins with 351 downregulated and 58 upregulated (**Fig. 4D, Table S4**). Pathway analysis of these proteins showed numerous strongly significantly enriched pathways, most notably with those related to glutamate receptors, synapse organization, and enzyme binding in the decreased proteins (**Fig. 4E,F**). A full list of all significantly enriched GO and KEGG pathways for both brain regions is included as **Tables S5-S8**.

Given the discrepancy in the number of proteins that were altered by G-CSF treatment during abstinence in the two brain regions, we also performed comparison analysis to see if this was an effect of magnitude of protein change, or if the cytokine treatment was indeed altering different pathways between the two regions. In **Figure 5**, we outline the most significantly altered pathways in each brain region and show overlap between the two. As is clearly seen, the regulation of proteins in the two brain regions is largely distinct other than some overlap in upregulated cytoplasmic proteins. These findings suggest that G-CSF during abstinence is having markedly different effects on the mPFC and NAc.

### G-CSF treatment during abstinence downregulates cytoskeleton, synapse formation, and glutamatergic signaling pathways in mPFC

To further interrogate the functional consequences of G-CSF treatment we performed STRING analysis to identify predicted protein-protein interactions among proteins altered by G-CSF treatment. Proteins downregulated by G-CSF in the mPFC form a densely interconnected network with 933 predicted protein-protein interactions which is significantly greater than the 550 that would be expected by chance (**Fig. 6A** – *p* < 1 × 10^−16^). Within this primary network there was an average of 5.44 interactions predicted per regulated protein. The upregulated proteins from mPFC, which were less abundant than those downregulated, also had significantly higher protein interactions than predicted by chance (36 edges, *p* = 0.0008, average node degree 1.26). However, differentially regulated proteins from NAc did not have significant increases in predicted interactions, again suggesting that the effects of G-CSF during abstinence seem to be driven by changes in mPFC. Full STRING statistics are available as **Table S9**.

**Figure 6.**
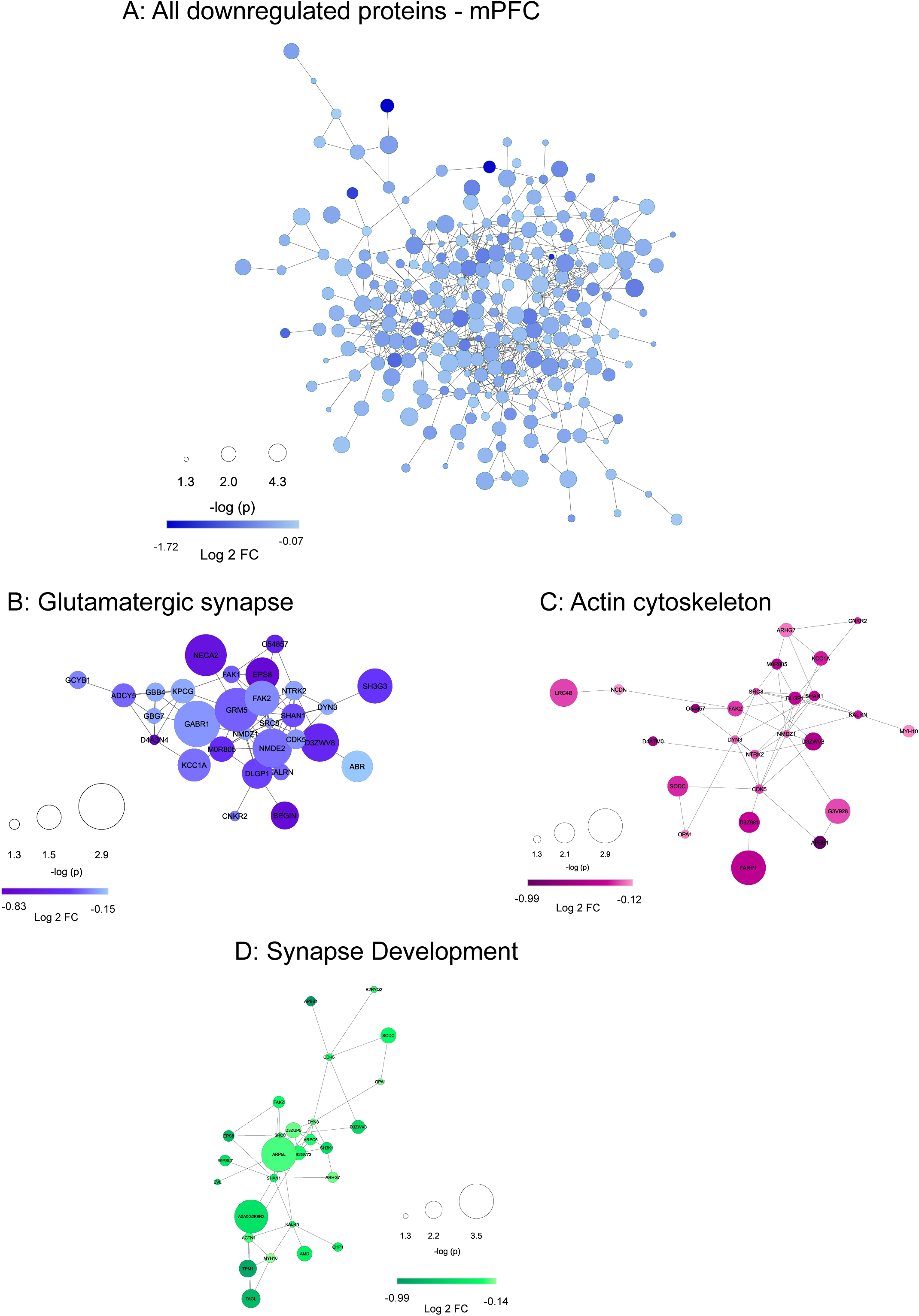
Downregulated proteins in the mPFC are from densely interactive networks with enrichment for synapses. (**A**) STRING protein interaction diagram of all proteins significantly downregulated by G-CSF in mPFC. Enrichment of interacting proteins is strongly statistically significant (*p* < 1.0 × 10^−16^) with an average number of 5.44 interactions per protein. Selected pathways were isolated from G:Profiler analyses to highlight interactions amongst downregulated proteins related to glutamatergic pathways (**B**), actin cytoskeleton regulation (**C**), and synapse development (**D**) related pathways – all of which have significantly more protein-protein interactions than would be predicted by chance (*p* < 1.0 × 10^−16^ for all). Protein nodes differ by size (-log p-value, larger node = smaller p-value) and color (darker color = greater fold change from PBS).

Within the downregulated mPFC proteins, we performed additional analyses on some of the most significantly regulated pathways related to synaptic function. Proteins involved in glutamatergic signaling were highly downregulated and create a densely interactive protein network (**Fig. 6B**). The same is true for proteins known to play key roles in regulation of the actin cytoskeleton (**Fig. 6C**) and synapse development (**Fig. 6D**). All of these pathways are critical for formation of appropriate structural and synaptic plasticity, and all form densely interconnected networks significantly above chance levels (*p* < 1 × 10^−16^ for all three). A merged version of these three networks is presented as **Figure 7** and shows how the proteins downregulated by G-CSF lead to decreases in signaling cascades involved in formation and potentiation of synapses in this critical brain region.

**Figure 7.**
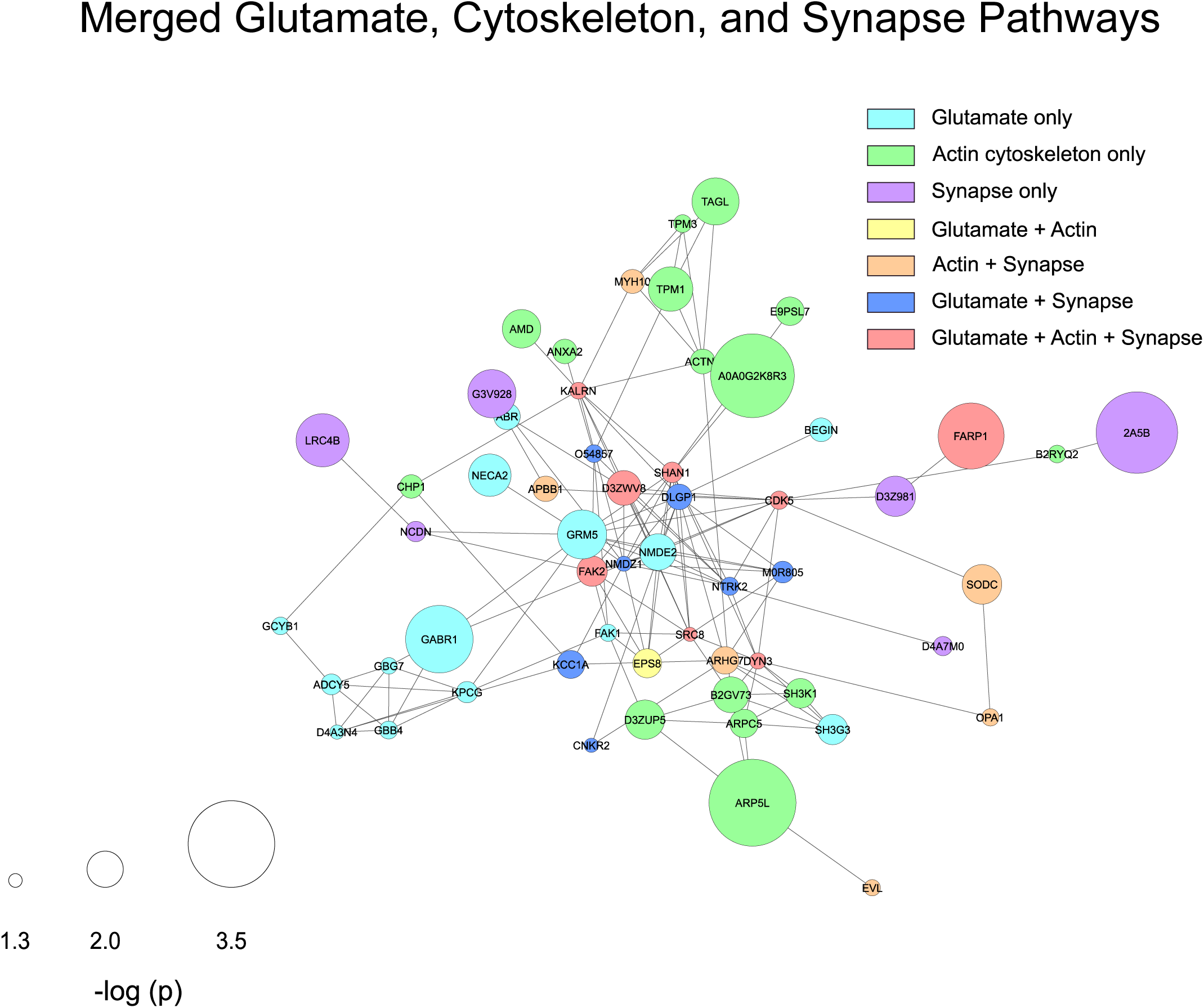
Overlap of synaptic and cytoskeletal protein interactions in the mPFC. Utilizing the regulated proteins from synaptic and cytoskeletal pathways, a STRING diagram was created showing the effect of G-CSF treatment on these critical regulators of synaptic plasticity and function. This network has 138 predicted protein-protein interactions compared to the 29 that would be predicted by chance (protein-protein interaction *p* value < 1.0 × 10^−16^; average node degree 3.83). Node color indicates pathway membership as shown in legend. Node size correlates with −log *p*-value compared to PBS control.

## Discussion

The current set of studies demonstrate that G-CSF administration accelerates extinction and reduces cue-induced cocaine-seeking (**Figs. 1 and 2**). While this seems contrary to the effects of G-CSF in potentiating effects on cocaine reward and self-administration (Calipari et al., 2018), there is a fair amount of evidence showing that G-CSF enhances learning in many different forms (Diederich et al., 2009a, 2009b; Kutlu et al., 2018). Integrating past findings with our current results, it seems G-CSF may be speeding response-outcome learning when administered during self-administration, and speeding learning when given during extinction or abstinence. Since extinction learning is a form of behavioral flexibility (Hamilton and Brigman, 2015), this study is also in line with Kutlu et. al demonstrating that G-CSF reduces trials to criterion in a reversal task (Kutlu et al., 2018). Interestingly, G-CSF did not influence cocaine-primed reinstatement. While both cues and a drug prime can reinstate cocaine-seeking, these types of reinstatement recruit unique but overlapping brain circuits (Kalivas and McFarland, 2003; Farrell et al., 2018). Alternatively, the injection of cocaine used for the prime might have interfered with the learning process.

This study also utilizes an unbiased approach to gain insight into the molecular consequences of G-CSF administration during abstinence that may underlie the reduced reinstatement. Proteomics analyses show that G-CSF causes significant changes in expression of ∼400 proteins in dorsal mPFC (**Fig. 4D**) but has less robust effects on protein expression in the NAc (∼40 proteins – **Fig. 4A**). We chose to examine these two brain regions as both the mPFC and NAc are necessary for cue-induced cocaine-seeking (Koya et al., 2009; Stefanik et al., 2013). Pathway analysis of the mPFC demonstrates significant changes in proteins related to glutamatergic signaling and synapse formation. Some examples of proteins downregulated include the scaffolding proteins Shank1 and Shank2, NMDA receptor subunits GluN1 and GluN2B, as well as mGluR5 and CaMKI (**Fig. 6, Table S4**).

Previous work has shown that pharmacological reduction of extracellular glutamate within the prelimbic cortex attenuates cocaine-seeking (Shin et al., 2018), highlighting the role of glutamate within this region in cue-induced reinstatement. Notably, the concomitant decreases in cue-induced reinstatement and GluN2B seen in our G-CSF treated rats are in line with previous literature. GluN2B is upregulated in mPFC during abstinence from extended cocaine access (Ary and Szumlinski, 2007; Ben-Shahar et al., 2009; Szumlinski et al., 2016) and inhibition of GluN2B-containing NMDA receptors in this area reduces cue-induced cocaine-seeking (Szumlinski et al., 2016). Additional work also confirms that mGluR5, another protein downregulated by G-CSF in the current dataset, is upregulated in dorsal mPFC 3 and 30 days after extended access cocaine self-administration (Ben-Shahar et al., 2013), although the behavioral relevance (if any) of mGluR5 in dorsal mPFC is unclear. G-CSF administration during abstinence might prevent upregulation of glutamatergic synaptic proteins normally altered by cocaine withdrawal in the mPFC.

In models of disease, G-CSF has repeatedly been shown to attenuate glutamate excitotoxicity, specifically in models of stroke (Schäbitz et al., 2003; Han et al., 2008) and Parkinson’s disease (Meuer et al., 2006). G-CSF administration (either systemic or intrathecally), reduces extracellular levels of glutamate (Han et al., 2008; Chen et al., 2010) via multiple mechanisms and has been shown to downregulate glutamate receptors (Diederich et al., 2009b; Mammele et al., 2016), an effect recapitulated in the current study. The elevated levels of extracellular glutamate that are present in the mPFC during abstinence from cocaine self-administration (Shin et al., 2016) likely do not cause excitotoxicity; however, G-CSF-induced downregulation of glutamate synaptic proteins might be regulating excitatory synaptic strength. While more work is necessary, this raises the intriguing possibility that G-CSF might scale glutamate activity to balance synaptic plasticity necessary for learning with a reduction in excitotoxicity produced by those same pathways.

Given that our previous work found that the NAc was the brain region responsible for the effects of G-CSF on cocaine place preference (Calipari et al., 2018), and that G-CSF increased stimulated dopamine release in the NAc (Kutlu et al., 2018; Brady et al., 2019), the disparity of the effects of G-CSF in the mPFC and the NAc was not expected. However, both of these regions receive synaptic inputs from the VTA which is also affected by repeated G-CSF treatment (Mervosh et al., 2018). Additionally, in our original studies we found that treatment with G-CSF potentiated cocaine induced c-fos activation in both the mPFC and the NAc, suggesting G-CSF exerts effects in both brain regions (Calipari et al., 2018). The differential effects we see here could reflect the difference between administration of G-CSF during a period of active cocaine administration vs. administration during abstinence, and may reflect the changes seen where G-CSF enhances cocaine intake during active administration (Calipari et al., 2018), but then hastens drug extinction and reduces cue-induced reinstatement when given during abstinence. It is important to note the timeline of the current experiments. Most work on incubation of cocaine craving compare neurobiology of reinstatement circuits after short-term cocaine withdrawal (1 and 3 days) to long-term or “incubated” cocaine withdrawal (28 to 30 days) (Grimm et al., 2001; Conrad et al., 2008). It is at this longer time point when neural changes first appear, such as insertion of GluR2-lacking AMPA receptors into NAc (Conrad et al., 2008). Since the current study employed a combination extinction / abstinence protocol that resulted in cocaine withdrawal for 18 days, it is possible that more protein changes would be observed in NAc at a later time point. In contrast with glutamatergic changes in the NAc, glutamatergic changes in mPFC occur quickly, with some effects appearing at 3 days (Huang et al., 2006) and most present by 14 days (Ben-Shahar et al., 2009).

Here we employed an unbiased proteomic analysis using data independent acquisition (DIA) to determine differences in protein expression between G-CSF and control treated rats during abstinence. The use of this technique is important for two reasons. The first is that by utilizing the DIA method, which bins proteins based on mass to charge (m/z) ratio (Wilson et al., 2019), we avoid the bias towards highly expressed proteins that is common with older data-dependent acquisition methods (Hu et al., 2016), and achieve more complete coverage of the proteome. Within these analyses we detected with confidence 2604 proteins within NAc expressed in all samples and 3364 in mPFC expressed in all samples (**Tables S1-S2**). The second is that by utilizing proteomics we can directly assess the functional state of the cell using a proteome-wide approach. Most studies in the field have used some version of transcriptomics to assess the molecular state of a cell, and while these methods are incredibly powerful and have revolutionized the field, levels of transcript are only important if they are translated into protein. Given the extensive regulatory mechanisms on mRNAs it would not be likely that there would be a strict 1:1 correlation of mRNA:protein (Payne, 2015), and indeed most studies that have examined these correlations find a correlation coefficient of *R*^*2*^ = 0.4 or lower (Ghazalpour et al., 2011; Schwanhäusser et al., 2011; Kumar et al., 2016). Thus, by directly querying protein changes induced by G-CSF treatment we can draw greater inference of how these treatments are altering the functional state of the cell.

Recombinant G-CSF is FDA approved for the treatment of neutropenia and is well-tolerated with a favorable side effect profile (Hoggatt and Pelus, 2014). Additionally, G-CSF has been tested in clinical trials for multiple neurological conditions including stroke and Alzheimer’s disease, thus establishing the feasibility of utilizing G-CSF as a therapeutic for a neuropsychiatric condition. This, along with the data presented herein, lay the foundation for a potential re-purposing of G-CSF for the treatment of SUDs. Importantly, this current work also identifies multiple protein targets for G-CSF’s effects in the mPFC and hints at a largely glutamatergic mechanism of action in its ability to reduce cocaine-seeking. The tolerability of G-CSF and uncovering of its cellular targets can guide the search for effective pharmacological interventions for psychostimulant use disorder.

## Supporting information

Supplemental Tables 1 through 9

## Acknowledgements

This work was funded by NIH grants DA-044308 & DA-049568 to DDK and DA-042111 & DA-048931 to ESC as well as by NARSAD Young Investigator Awards to RSH, ESC & DDK. The Orbitrap Fusion mass spectrometer and the Offline UPLC utilized were supported in part by NIH SIG grants 1S10OD019967-0 & 1S10ODOD018034-01, respectively, and Yale School of Medicine. This work was supported by the Yale/NIDA Neuroproteomics Centre Grant DA-018343. Cocaine was provided by the NIDA drug supply program.

## Disclosures

All authors declare no financial conflicts of interest.

