## Supplementary figures and images for "Granulocyte-Colony Stimulating Factor reduces cocaine-seeking and downregulates glutamatergic synaptic proteins in medial prefrontal cortex"

### Supplemental figure 1

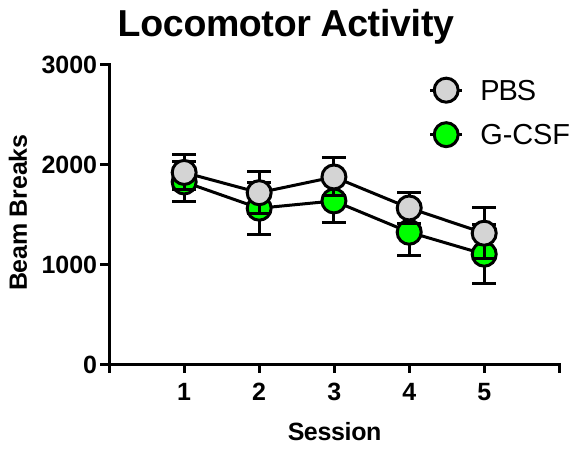

**Figure S1**
